# Evidence of MHC class I and II influencing viral and helminth infection via the microbiome in a non-human primate

**DOI:** 10.1101/2021.06.01.446545

**Authors:** B. Karina Montero, Wasimuddin, Nina Schwensow, Mark A. F. Gillingham, Yedidya R. Ratovonamana, S. Jacques Rakotondranary, Victor Corman, Christian Drosten, Jörg U. Ganzhorn, Simone Sommer

## Abstract

Until recently, the study of major histocompability complex (MHC) mediated immunity has focused on the direct link between MHC variability and susceptibility to parasite infection. However, MHC genes can also influence host health indirectly through the sculpting of the bacterial community that in turn shape immune responses. We investigated the links between MHC class I and II gene variability gut microbiome diversity and micro- (adenovirus, AdV) and macro- (helminth) parasite infection probabilities in a wild population of non-human primates, mouse lemurs of Madagascar. This setup encompasses a plethora of underlying interactions between parasites, microbes and adaptive immunity in natural populations. Both MHC classes explained shifts in microbiome composition and the effect was driven by a few select microbial taxa. Among them were three taxa (*Odoribacter*, *Campylobacter* and Prevotellaceae-UCG-001) which were in turn linked to AdV and helminth infection status, evidence of the indirect effect of the MHC via the microbiome. Our study provides support for the coupled role of MHC variability and microbial flora as contributing factors of parasite infection.

**Author summary:** The selective pressure of the major histocompatibility complex (MHC) on microbial communities, and the potential role of this interaction in driving parasite resistance has been largely neglected. Using a natural population of the primate *Microcebus griseorufus*, we provide evidence of two outstanding findings: that MHCI and MHCII diversity shapes the composition of the gut microbiota; and that select taxa associated with MHC variability predicted adenovirus and helminth infection status. Our study highlights the importance of incorporating the microbiome when investigating parasite-mediated MHC selection.

## Introduction

Reciprocal interactions between the hosts’ immune system and the gut microbial flora are essential in defining the course of a parasitic challenge for three main reasons. First, specific bacterial taxa can prime immune signaling that provoke the activation of an inflammatory response against parasites [1, 2]. Second, the secretion of inhibitory substances (e.g. antimicrobials) and metabolites by some bacteria have been shown to inhibit growth and the ability of pathogens to attach to the intestinal lumina [3, 4]. Third, the competitive advantage of commensal bacterial taxa over a niche can limit the growth and expansion of pathogenic bacteria [5, 6]. Evidence of these and other forms of colonization resistance underscore the relevance of immune mechanisms that encourage a balanced state between tolerance towards commensal bacteria and resistance against offending invaders [7, 8]. An important factor influencing this balance is the selective sculpting of bacterial communities by means of adaptive immunity [9–11]. The involvement of the major histocompatibility complex (MHC) in antigen-specific immune response appears to contribute to microbial colonization [12–14], however the interplay between the natural variation of the MHC and microbial communities in parasite resistance has yet to be explored.

Antigen-specific responses are mediated by two distinct classical pathways contingent of the nature of the foreign peptide; MHC class I molecules process intracellular peptides (e.g. viruses and cancer cells) and present them to CD8^+^ T-cells while extracellular peptides (e.g. bacteria and helminths) are presented to CD4^+^ T-cells by MHC class II molecules [15, 16]. The high affinity of CD4^+^ T-cells to the bacterial antigen/MHC class II complex trigger inflammatory [1, 17, 18] and tolerogenic [19] responses, which in turn keep commensals in check. Therefore, most studies focus on the role that the MHC class II pathway plays in the gut microbiome. Evidence of compositional differences in the microbiome associated to primed CD8^+^ T-cells by MHC class I molecules is scarce, although the MHC class I pathway synergizes with specific bacterial taxa to mediate parasite resistance [2, 20] and tumor development [21]. Furthermore, processing of extracellular antigens through the specialized role of dendritic cells for cross-presentation [22] has been associated with potent T-cell responses [23]. Therefore, both CD4^+^ and CD8^+^ T-cells are likely involved in tolerogenic or immune responses towards parasites and gut commensals.

Here, we used a natural population of the reddish-gray mouse lemur *Microcebus griseorufus* to reconcile the inter-dependent relationship of MHC class I, class II and microbial diversity on micro- and macro- parasite infection propensity (a detailed summary of the study design and hypotheses are given in Figure 1 (Fig 1). To this end, we explored adenovirus (AdV) and helminth infection probabilities. AdV is a DNA virus causing widespread infections and are considered a major concern for human health globally [24]. Although AdV infection is often asymptomatic, it can develop into severe respiratory and gastrointestinal diseases in children, immunosuppressed patients and in the elderly [25]. It is also important to monitor AdV prevalence in wildlife, in particular in non-human primates, since interspecies transmission are known to occur and can impact human health [26, 27]. In contrast, helminths play important immunomodulatory roles in the establishment of long-lasting infections that can be either harmful, particularly for undernourished and immunosuppressed individuals [5], or beneficial to the host by limiting the development of immune pathology, a process that involves the concerted action with the microbiota [28, 29]. Chronic viral infection, helminths and commensal bacteria share the common agenda of promoting acceptance by its host, often via immune regulation. Our group previously demonstrated a link between AdV and gut microbiome using 16SrRNA gene amplicon data [30]. Given its central role in immunity, we expand on our previous findings and tested the hypothesis that diversity at the MHC is an important component of the host’s genetic landscape in shaping gut microbial communities, which in turn influence micro- and macro- parasite infection.

**Fig 1.**
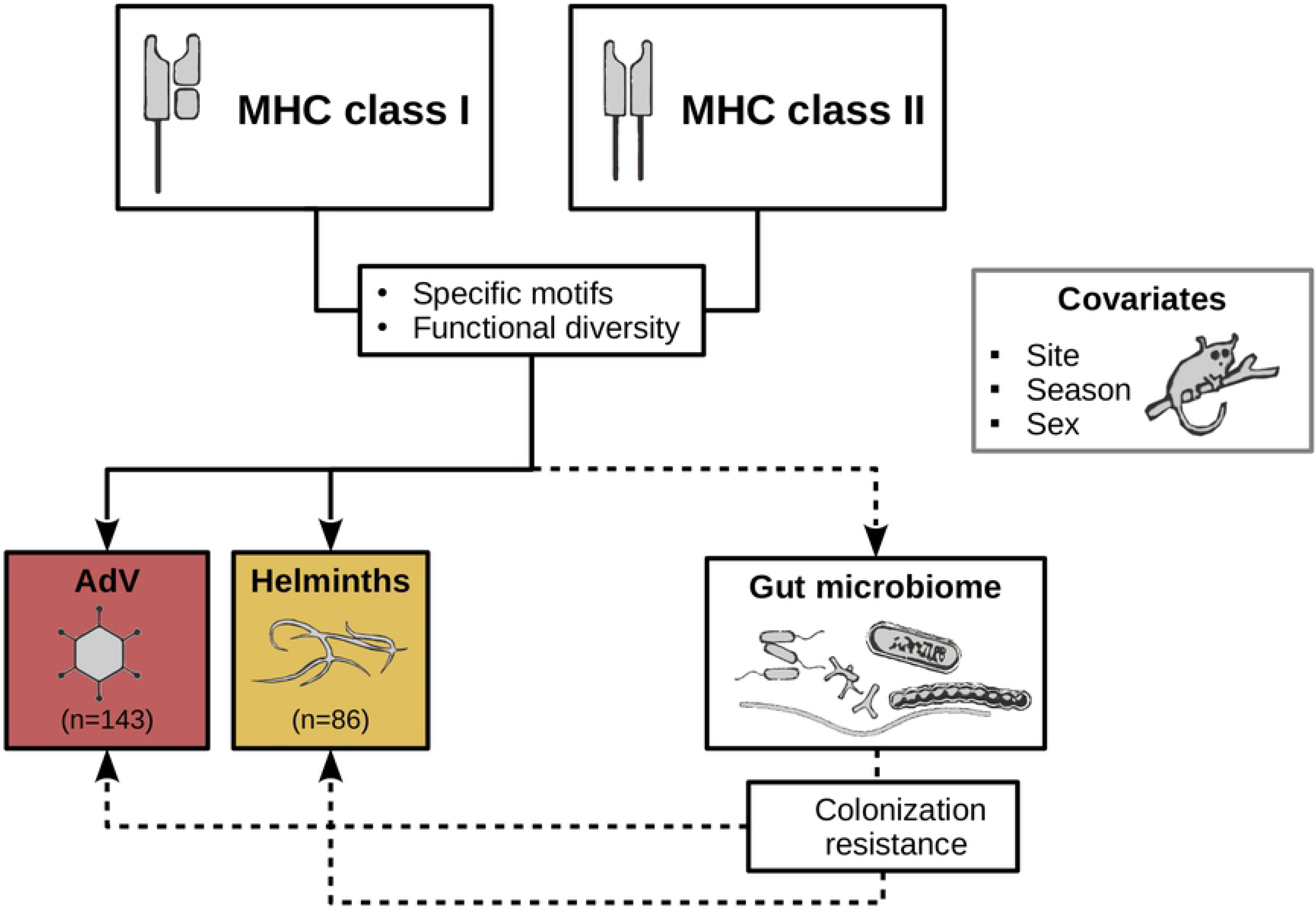
Study design. The role that MHC polymorphisms may play on parasite infection in natural populations can be better understood under the lens of their direct and indirect contribution. The MHC may have direct effects on parasite infection via antigen-driven immune response. It may also indirectly affect the environment that parasites face when establishing an infection by shaping the gut microbiome. We evaluated the direct (solid arrows) and indirect (dashed arrows) contribution that MHC class I and class II variability may have on micro- (adenovirus, AdV) and macro-parasite (helminths) infection status in a wild population of non-human primates, the mouse lemur *M. griseorufus*, while controlling for the effect of extrinsic (sampling site and season) and intrinsic (sex) factors. We assessed MHC variability in terms of the presence of specific motifs (MHC class I supertypes and MHC class II alleles) and functional diversity (sequence divergence and number of supertypes). We evaluated the direct effects of the MHC on infection status according to the following predictions: i) specific MHC class I supertypes and MHC class II alleles will be associated with susceptibility (or resistance) to AdV and helminth infection, respectively; ii) functional diversity will be negatively associated with parasite infection status. For the indirect effects of the MHC on infection status via the microbiome, we predicted that MHC I and MHC II variability influences the diversity and the composition of the microbiome. In particular we predicted that specific MHC motifs and MHC functional diversity will be negatively associated with the relative abundance of pathogenic taxa. In contrast, we expect to find that taxa not subject to immune recognition mediated by the MHC are involved in essential metabolic functions and contribute to resistance to parasitic challenges.

## Results

### MHC motifs and the microbiome predict infection status

Consistent with previous work on mouse lemurs [27, 31], AdV and helminth prevalence in our study population was high; 29.4% of individuals tested AdV^+^ (n=143), 40.1% of individuals (n=86) were infected with helminths, and 8.6% of individuals were co-infected with AdV and helminths. We detected a total of 226 MHCI functional alleles (median of 7 alleles per individual, range 4-20) in the population and grouped them into 17 MHC supertypes (ST) based on physicochemical variables (z-values [32]) of the amino acid sites affected by positive selection (PSS) (*Materials and Methods*, S1 Fig and S1 Table). For the MHCII, we identified 49 functional alleles. A large number of MHCII alleles were rare (present in 1-2 individuals), therefore we used for subsequent analyses only alleles that were found in more than three individuals (S2 Fig). We found that two MHC motifs were associated with AdV infection status (MHCI ST^*^3 and MHCII Migr-DRB^*^34 allele)(S3 Fig C and E). However, neither MHCI nor DRB motifs were associated with helminth infection status (S3 Fig D and F).

We used binomial GLMs to examine the effect of MHC and microbiome diversity on parasite infection while controlling for the potential effects of covariates (site, season and sex). We performed model selection using the information theoretic (IT) approach and we estimated the effect of predictors using weighted model averaging (see *Materials and Methods*) [33]. We used the number of MHCI supertypes (MHCI_nST_), mean amino acid allele divergence over PSS at MHCI gene (MHCI_distPSS_), and mean amino acid allele divergence over PSS at MHCII gene (MHCII_distPSS_) as proxies of MHC functional diversity (*Materials and Methods*). Additionally, the presence of MHC motifs associated with AdV infection (ST*3 and Migr-DRB*34, S3 Fig) was included in the binomial models in order to test for the relative contribution of specific motifs vs functional diversity on AdV infection status. For the microbiome we distinguished between diversity (Faith’s PD and Shannon’s index of diversity) and divergence (Bray-Curtis distance from the population mean). The former quantifies gut microbial diversity within an individual, whilst the latter quantifies how atypical a host’s microbiome composition is relative to the population mean.

We found strong support for the contribution of a single MHC motif on AdV infection status. Individuals carrying the MHCII DRB allele Migr-DRB*34 are more likely to be AdV^pos^ (Δ AIC= 3.04; Cohen’s D [±95%CI] = 0.84 [0.06,1.60])(Fig 2C). However, we found no support of an association between MHC functional diversity and AdV or helminth infection status according to model selection (S2 Table - S7 Table), suggesting that a specific MHC motif plays a primary role in AdV infection status relative to MHC functional diversity. In addition, we found that microbiome diversity and divergence were associated with AdV and helminth infection, respectively. AdV infection was associated with an increase in Faith’s PD (Δ AIC= 6.06; partial-r [±95%CI] = 0.26 [0.15, 0.37]) (Fig 2B, S2 Table and S6 Table). We found a negative relationship between microbiome divergence and helminth infection status (Δ AIC= 2.26; partial-r [±95%CI] = −0.22 [−0.41,−0.04]) (Fig 2D, S3 Table and S6 Table), indicating that gut microbial communities across helminth infected individuals are more similar than the communities of non-infected individuals. A relationship between Shannon’s diversity index and AdV or helminth infection probability was not supported by model selection (S4 Table, S5 Table and S7 Table). With respect to the influence of other covariates on parasite infection, we found strong support for the effect of season on helminth infection status (Δ AIC= 5.19; Cohen’s D [±95%CI] = −0.49 [−0.98,−0.02]) (Fig 2E, S3 Table and S6 Table); the probability of infection was higher among individuals captured during the dry season, as has been demonstrated in baboons [34]. Limited food availability during the dry season relative to the wet season results in nutritional stress which is considered an important driver of increased parasite burden [35].

**Fig 2.**
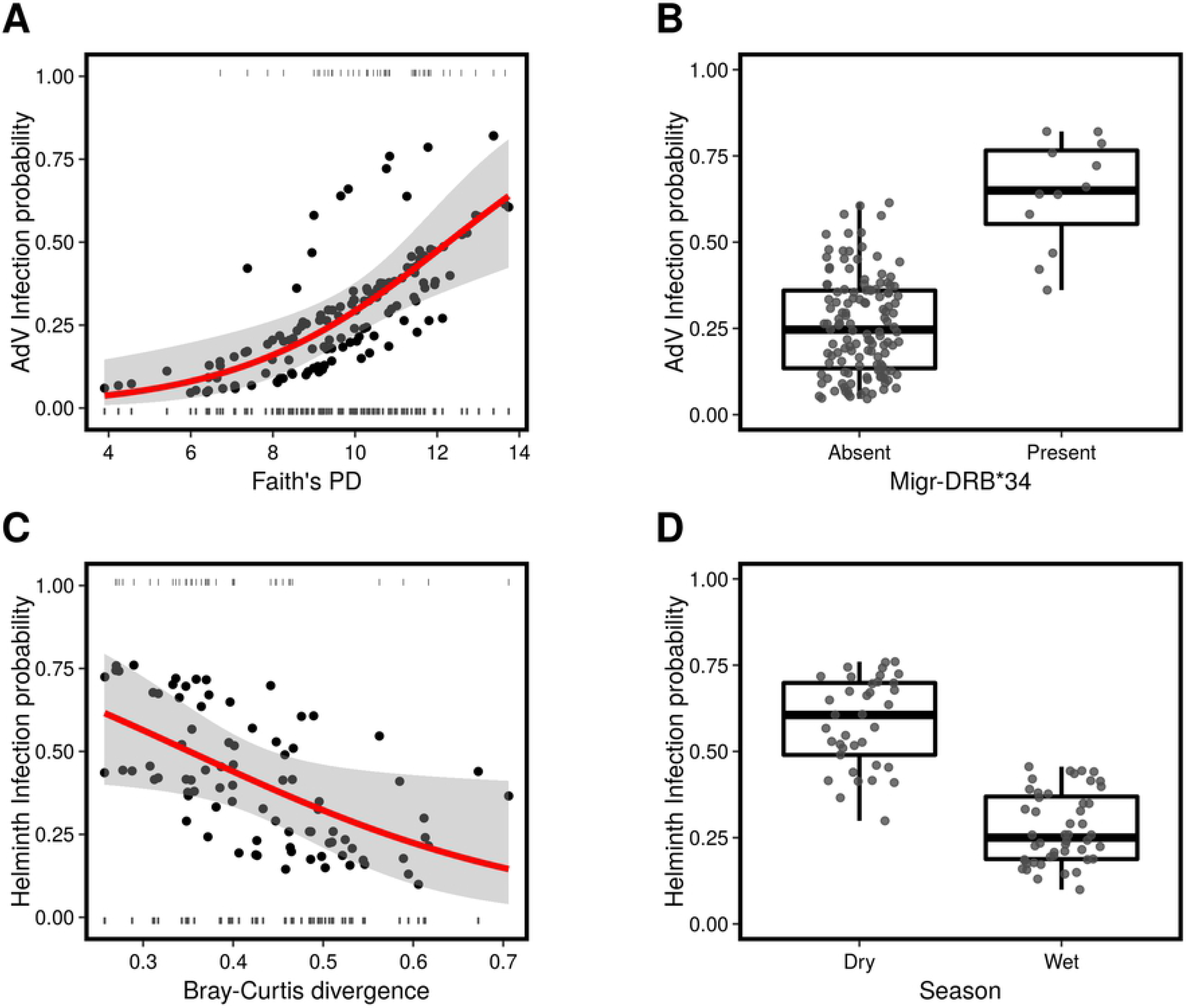
Association between MHC functional diversity, microbiome diversity (Faith’s PD) and AdV and helminth infection. Faiths’ PD and the presence/absence of Migr-DRB*34 were retained as predictors of AdV infection probability by model selection, whilst Bray-Curtis divergence and season were retained as predictors of helminth infection probability. AdV infection probability according to model averaged predicted values of *(A)* Faiths’ PD and *(B)* the presence/absence of Migr-DRB*34. Helminth infection probability according to model averaged predicted values of *(C)* Bray-Curtis divergence and *(D)* season. Fitted lines in *(A)* and *(C)* are shown in red and 95% confidence intervals are shaded in gray.

### MHC variability is associated with a shift in the composition of the gut microbial community

To investigate the indirect role of the MHC on parasite infection via the microbiome, we explored the association between MHC variability and microbiome diversity and composition. Using regression models we found no evidence that MHC diversity is associated with microbial diversity within an individual (S8 Table). In contrast, gut microbiome composition was associated with MHC variability. We used canonical correspondence analysis (CCA) analysis to explore the contribution of MHC functional divergence, specific MHC motifs and covariates shaping the composition of the gut microbial community. Our results revealed that sequence divergence at MHCI_distPSS_ (r^2^=0.16, *P*<0.0001) (but not MHCI_nST_) was a significant predictor of distance between samples in the constrained ordination space (S4 Fig). Contrary to expectations based on the affinity of class I vs class II to extracellular peptides, we did not find evidence of an effect of MHCII variability on microbial composition (r^2^=0.03, *P* =0.09). We observed shifts in microbial composition according to the presence of eight MHCI supertypes (S5 Fig) and four MHCII alleles (S6 Fig). Regarding the effect of parasite infection on composition, both AdV infection status (r^2^=0.03, *P* =0.01; S7 Fig A) and helminth infection predicted a shift in microbial composition (r^2^=0.05, *P* =0.02; S7 Fig B). The former result replicates our previous findings that AdV^pos^ individuals have a different microbiome from AdV^neg^ individuals [30]. Ordination analyses also reveal a significant effect of season (r^2^=0.37, *P*<0.0005; S7 Fig C) and sampling site (r^2^=0.21, *P*<0.0005; S7 Fig D), highlighting the fine-scale temporal and spatial patterns of microbial diversity [36]. Sex had no effect on beta diversity (r^2^=0.005, *P* =0.47). Our study sites differ sharply in vegetation mostly as a consequence of differences in precipitation regimes. The strong effects of site and seasonal on gut microbiome composition are likely to reflect differences in food availability and dietary preferences [37, 38].

We identified core taxa (prevalence > 60%) associated with specific MHCI supertypes and MHCII alleles (using analysis of composition of microbes (ANCOM)) and that differed most with respect to their association with high or low estimates of MHC functional diversity (using a differential ranking approach) (see *Materials and Methods*). We found that both MHCI and MHCII variability were linked to shifts in the relative abundance of a limited number of core taxa (Fig 3). In fact, except for one MHCII allele (Migr-DRB*36) which was associated with seven core ASVs (and with 22 ASVs in total) and was strongly associated with an overall shift in microbial composition (r^2^=0.16, *P*<0.001; S6 Fig C), the vast majority of MHCI supertypes and MHCII alleles were associated with a small range (1-3) of ASVs (Fig 3A and B). This suggests that Migr-DRB*36 is a generalist [39] that indiscriminately binds a high number of antigens affecting both commensal and potentially pathogenic taxa.

**Fig 3.**
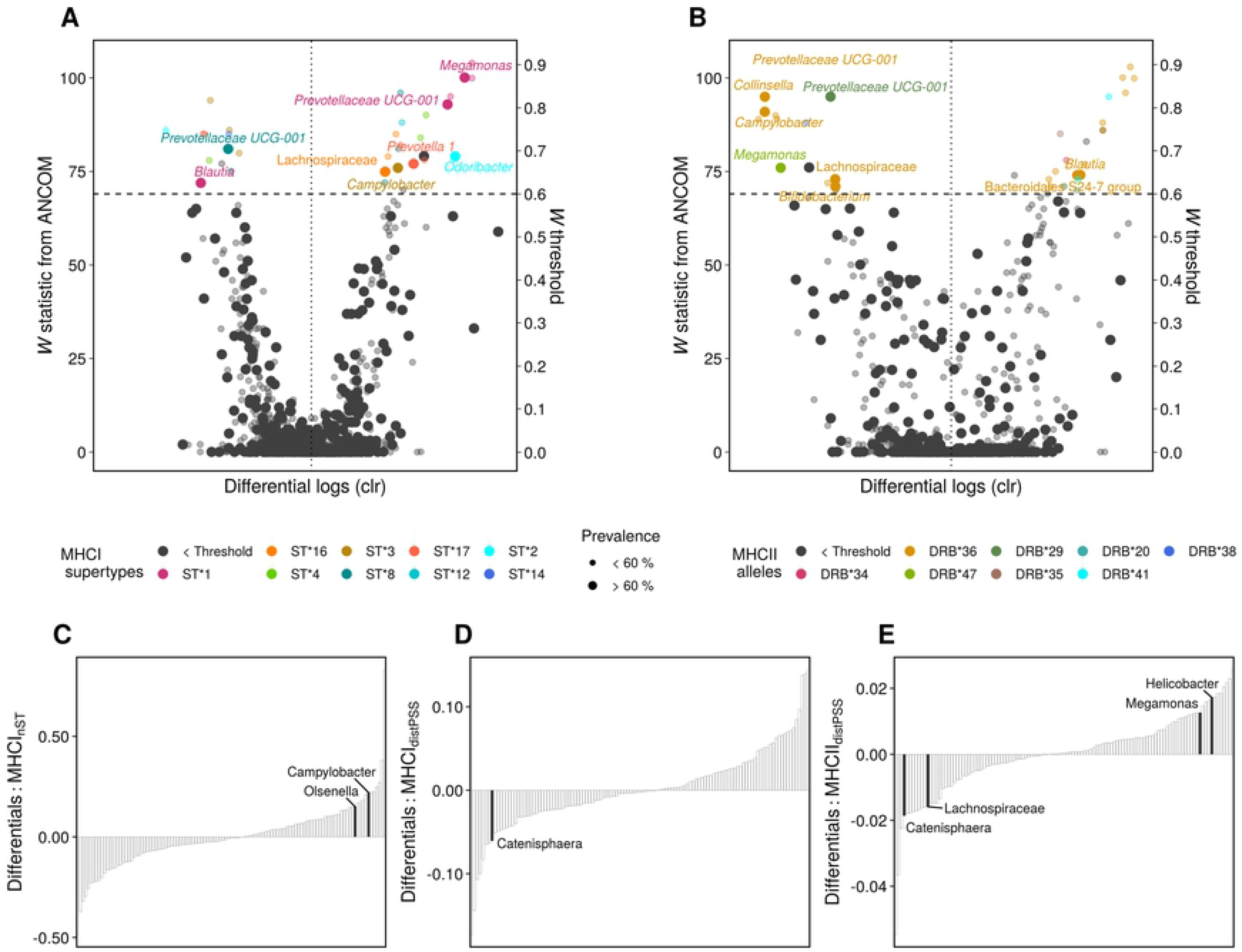
MHC variability is associated with shifts in microbiome composition. *(A-B)* Association between specific MHCI supertypes and MHCII alleles and microbial taxa (n=143). Significance threshold is informed by W statistic (horizontal dashed line) and F estimates (vertical dotted line). ASV’s above this threshold represent taxa significantly (*P* < 0.005) associated with a specific MHC motif. Taxa are colored according to their association with a MHC motif. Labels are shown for core taxa (prevalence >60%). *(C-E)* Microbial ranks based on multinomial regression coefficients sorted by their association with supertype diversity (MHCI_nST_), MHCI sequence divergence (MHCI_distPSS_) and MHCII sequence divergence (MHCII_distPSS_) (n=143). Labels correspond to high and low rankings of core taxa.

### Evidence of an indirect effect of the MHC on parasite resistance

We then used log-ratios, a suitable method for examining differences in abundance using compositional data [40], to estimate differences in relative abundance of the ASV’s associated to MHC variability (MHC motifs and functional diversity) according to AdV and helminth infection status. We used *Bifidobacterium* as a reference frame since this taxa was present in most samples, and is expected to be a stable member of the microbial community (*Materials and Methods*). Our analyses indicate that resistance to AdV was provided by an increase in *Campylobacter* (Fig 4 B) which was associated with both MHCI and MHCII diversity (MHCI supertypes: ST*3, ST*4; MHCII allele Migr-DRB*36 and MHCI_nST_, Fig 4 A, B and C), an increase in *Olsenella* (Fig 4 C), a microbe thought to prevent inflammation [41] associated to MHCI_nST_ (Fig 4 C), and an increase in abundance of *Odoribacter* (Fig 4 D), a microbe enriched among individuals with the MHCI supertype ST*2 (Fig 4 A). Furthermore, the ASV assigned to the Prevotellaceae UCG-001 group and that was associated to several MHC motifs (MHCI supertypes: ST*1 and ST*8; MHCII alleles: Migr-DRB*36 and Migr-DRB*29) was linked to helminth infection (Fig 4E).

**Fig 4.**
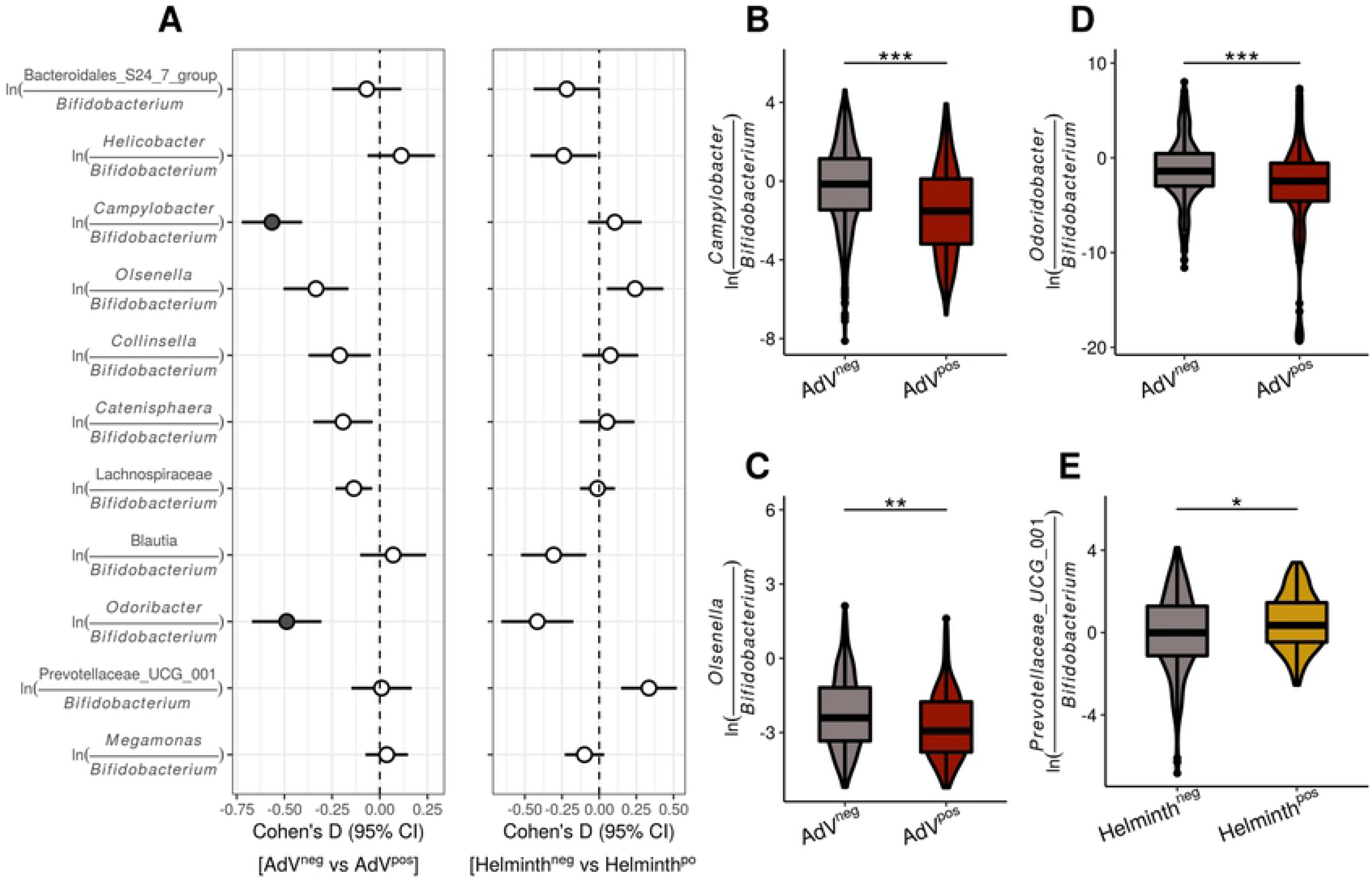
Bacterial taxa associated with MHC motifs and functional diversity are significantly associated with AdV and helminth infection status. *(A)* Effect size forest plots for the effect (Cohen’s D and CI) of AdV and helminth infection on the log-ratios between taxa associated to MHC motifs and to MHC functional diversity. Filled symbols represent log-ratios with medium to large effect sizes (Cohen’s D |>0.50|). ASVs of the genus *Bifidobacterium* were used as a reference frame to calculate the log-ratios. *(B-E)* Violin box-plots illustrating the significant log-ratio across AdV (n=143) and helminth (n=86) infection status. * *P* < 0.05; ** *P* < 0.005; *** *P* < 0.0005.

Overall, more taxa associated with MHC diversity were also linked to AdV resistance compared to the number of taxa influencing helminth infection, suggesting a stronger indirect effect of the MHC on AdV infection propensity than helminth infection status. Notably, among the taxa associated with infection, both *Campylobacter* and *Odoribacter* had a particularly strong effect on AdV propensity (Fig 4 A). *Campylobacter* is a genus well known for its high number of important pathogens for human and animal health [42], whilst both *Odoribacter* and Prevotellaceae UCG-001 are known to generate short-chain fatty acids (SCFA) [43, 44].

### Links between MHC variability and diversity of functional bacterial pathways

To evaluate if the observed shift in taxonomic composition of the microbiome influences functional attributes, we generated and analyzed the abundance of MetaCyc pathway predictions using the PICRUSt2 algorithm [45]. MetaCyc is an open-source database for metabolic pathways based on organismal annotated genomes [46]. We first evaluated whether MHC functional diversity and covariates predicts pathway diversity (pathway richness and evenness using Shannon’s index) and composition. We included a total of 296 MetCyc pathways in the analysis. Greater peptide recognition among individuals with high levels of MHC functional diversity can constrain the ability of a variety of bacterial taxa to colonize the gut, a scenario that would be supported by a negative linear coefficient in functional microbial diversity. In contrast, a positive linear coefficient would suggest that maximal MHC diversity is associated with the maintenance of high functional microbial diversity as a result of the removal of specific taxa that negatively impact other members of the microbial community. We found that MHCII sequence divergence was negatively associated with microbiome pathway evenness (Δ AIC= 2.26; partial-r [±95%CI] = −0.22 [−0.41,−0.04]) (Fig 5 C), S9 Table), suggesting that MHCII functional divergence limits gut microbial functional diversity. The fact that MHCII sequence divergence was associated with pathway evenness (Shannon’s diversity Index) but not pathway richness, suggests that the effect of the MHC was stronger on abundant core pathways than on sparse pathways. None of the MHCI estimates were associated with pathway diversity (S9 Table), suggesting that MHCI molecules do not influence the overall functional diversity of the microbial community. In contrast to taxonomic based analyses, AdV and helminth infection status had no effect on pathway diversity (S9 Table and S10 Table). With respect the effect of covariates, we found support for temporal variation in pathway eveness; pathway eveness in the second dry season of sampling was higher than in all other sampled seasons (Δ AIC= 15.9; Cohen’s D _Dry2 vs Dry1_ [±95%CI] = −0.70 [−1.08,−0.31], Cohen’s D _Dry2 vs Wet1_ [±95%CI] = −0.77 [−1.17,−0.38], Cohen’s D _Dry2 vs Wet2_ [±95%CI] = −0.51 [−0.88,−0.13], S9 Fig A). In addition, we found an effect of sampling site on pathway richness (Δ AIC= 7.02; Cohen’s D [±95%CI] = −0.58 [−1.00,−0.14], S9 Fig B), further emphasising previously mentioned habitat specific effects on the gut microbiome.

**Fig 5.**
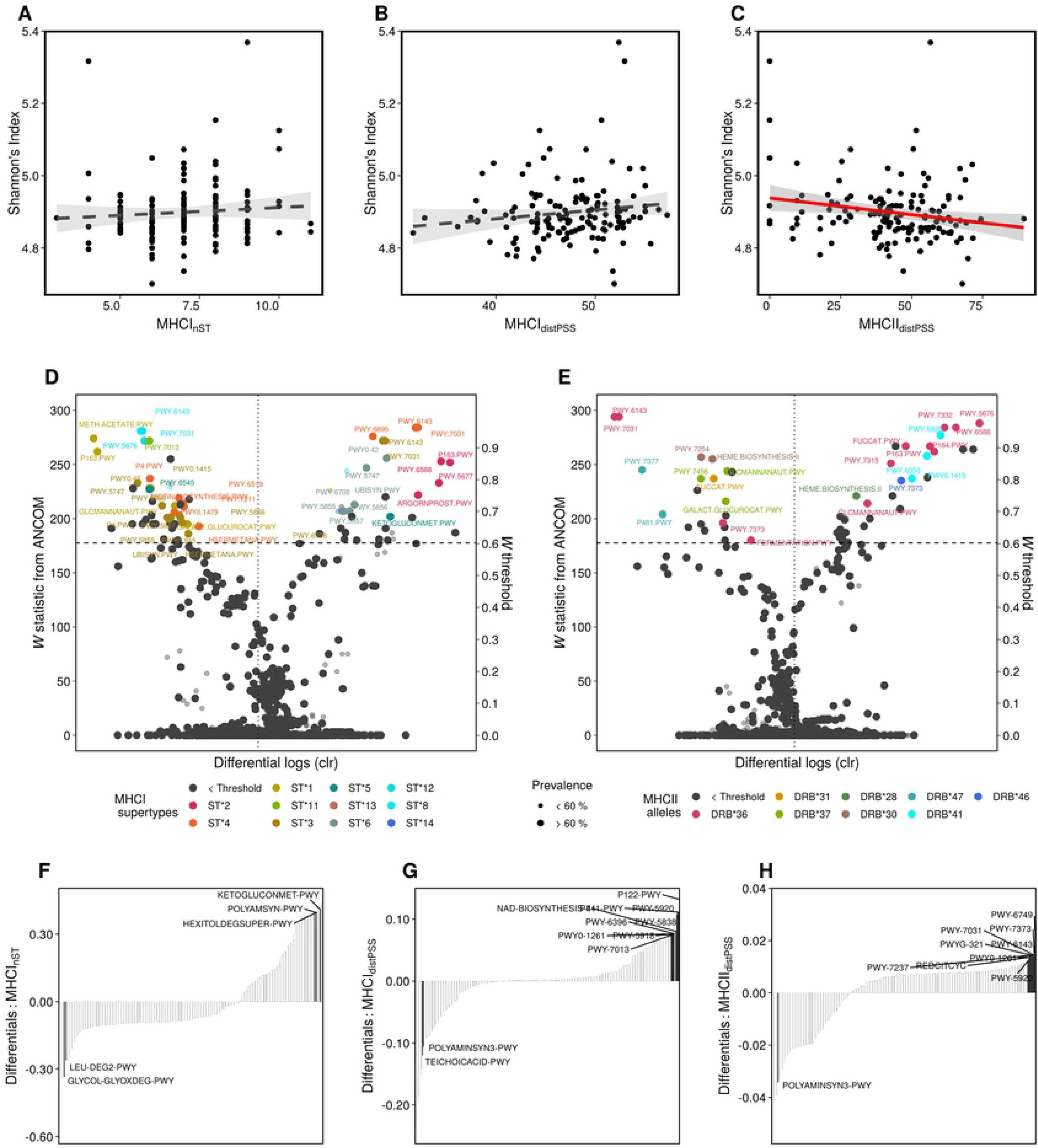
MHC variability is associated with diversity and composition of functional pathways. Relationship between *(A)* number of MHCI supertypes, *(B)* MHCI sequence divergence and *(C)* MHCII sequence divergence and the functional diversity of the microbiome according to Shannon’s diversity index. Associations supported by model selection (Δ AIC > 2) are reported with a red solid line, shaded area represent 95% confidence intervals. *(D-E)* Association between specific MHCI supertypes and MHCII alleles and metabolic pathways. Significance threshold is informed by W statistic (horizontal dashed line) and F estimates (vertical dotted line). Metabolic pathways above this threshold represent pathways significantly (*P* < 0.005) associated with a specific MHC motif. Pathways are colored according to their association with a MHC motif. *(F-H)* Microbial pathway ranks based on multinomial regression coefficients sorted by their association with supertype diversity (MHCI_nST_), MHCI sequence divergence (MHCI_distPSS_) and MHCII sequence divergence (MHCII_distPSS_). Labels correspond to high and low rankings of core pathways (>60% prevalence).

CCA analysis indicate that the MHC had limited effects on the overall composition of metabolic pathways. On the one hand, we found fewer associations between specific MHC motifs (2 MHCI supertypes and 1 MHC II allele, S8 Fig) and among-individual distance in the ordination space compared to taxonomic-based analyses. On the other hand, none of the MHC metrics of functional diversity were significant predictors of the composition of pathways. Furthermore, consistent with taxonomic based analyses we also detected significant temporal variation (r^2^=0.15, *P*<0.0005) in the composition of functional pathways (S9 Fig C)).

As a next step we identified pathways whose relative abundance was associated to specific MHC motifs and pathways with high and low ranks according to MHC functional diversity and then used log-ratios to test for differences among AdV and helminth infected vs non-infected individuals. To this end, we used ANCOM and differential ranking as described above for taxonomic-based analyses. We found that a particular set of MHC motifs were linked to differences in the relative abundance of a restricted number of pathways (Fig 5 D-E). Mirroring our results on bacterial taxa, most motifs were associated with a few pathways (1-4), with the exception of ST*3 and Migr-DRB*36 which were associated with differences in the relative abundance of a larger number of pathways (14 and 12 pathways, respectively). Finally, multinomial analyses revealed that low MHCI and MHCII functional diversity (for both number of supertypes and sequence divergence) was associated with relatively rare pathways, while core pathways were associated with high MHCI and MHCII sequence divergence (Fig 5 F-H).

To calculate log-ratios we used the TCA pathway (PWY-7254) as a reference since this is a common pathway central to the generation of energy and present in most aerobic living organisms. We did not find significant differences in the relative abundance of pathways according to parasite infection. However for this analysis we had large effect size confidence intervals (as a result of strong inter-individual stochasticity in pathway abundance) suggesting a lack of statistical power. For instance, when considering pathways with medium to large effect sizes (| Cohen’s D| > 0.5), we identified three pathways with a lower relative abundance among individuals testing positive for AdV and helminth infection (Fig 6 A). The relative abundance_relPWY-7254_ of pathways involved in the generation of glycan (PWY-7031) and sugar (PTW-6143) were increased among AdV^neg^ individuals (Fig 6 A), suggesting that these pathways are involved in limiting AdV infection. Interestingly, the stratified pathways revealed that *Campylobacter* contributes entirely to the diversity of these pathways (Fig 6 B-D). Indeed, PWY-7031 and PTW-6143 have been associated with adherence and pathogenicity, which suggests that *Campylobacter*-enriched functional pathways associated with high MHCII sequence divergence and multiple MHC motifs (MHCI supertypes: ST*1, ST*4 and S*8; MHCII allele Migr-DRB*36) reduce AdV susceptibility. A single metabolic pathway (PWY-5920), decreased in abundance among helminth^neg^ individuals (Fig 6 D). This pathway is involved in heme biosynthesis, an iron prosthetic group that is essential in the development and reproduction of helminths [47].

**Fig 6.**
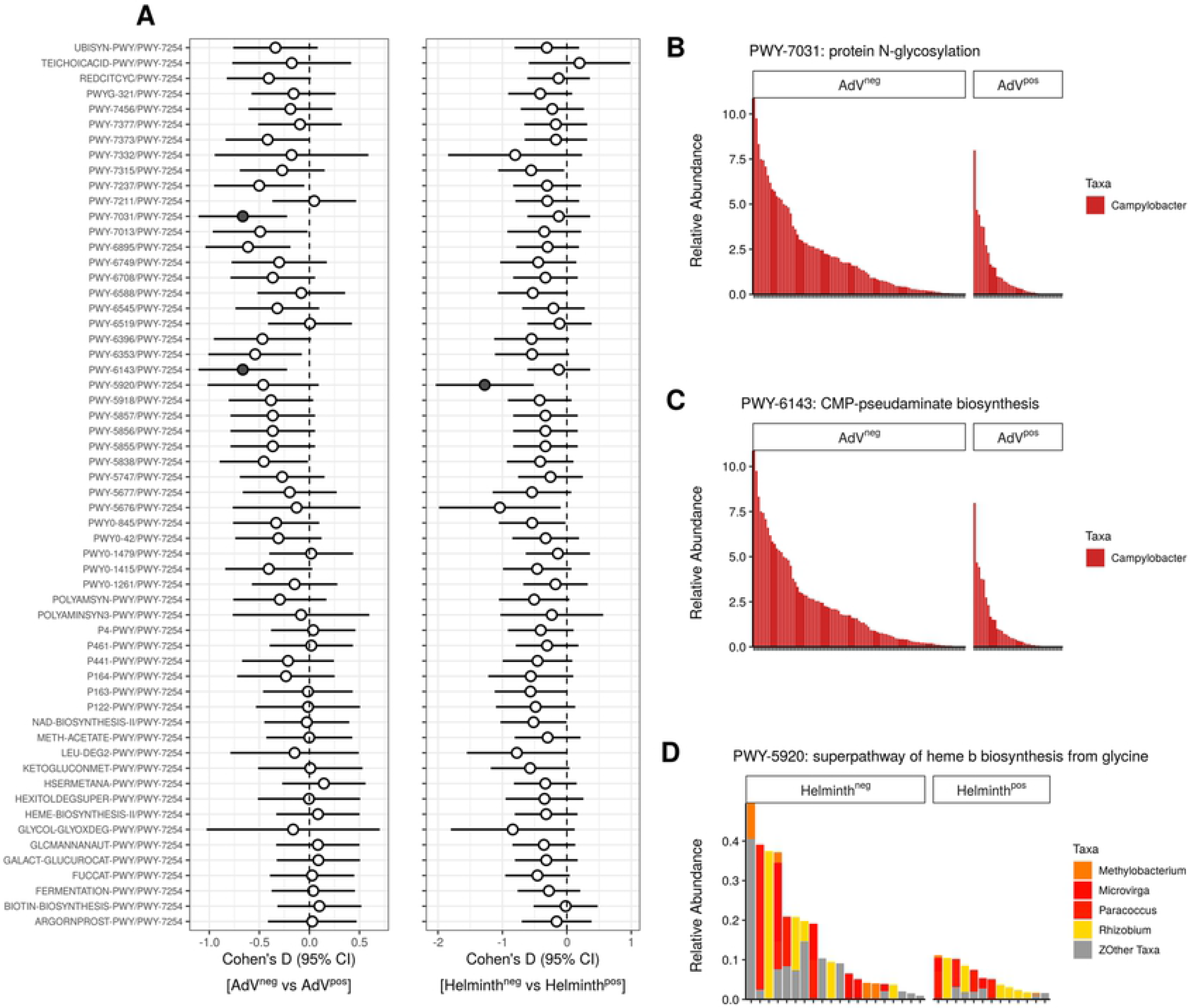
Differences in relative abundance of functional pathways associated with MHC variability according to AdV and helminth infection. *(A)* Effect size forest plots of the log-ratios of the functional pathways identified by ANCOM and differential ranking with respect to AdV and helminth infection. The pathway PWY-7254 was used as a reference frame to calculate the log-ratios. Filled symbols represent log- ratios with medium to large effect sizes (| Cohen’s D| > 0.50). *(B-D)* Contribution of bacterial taxa to pathways with medium-large effect sizes according to with AdV or helminth infection. Stacked bar plots indicate the relative abundance of taxa across samples.

## Discussion

The contribution of MHC variability on parasite resistance has traditionally been evaluated as a direct interaction. Indeed, parasite mediated selection is considered the main mechanism driving the extraordinary polymorphism of MHC genes [48, 49]. However, parasites do not only face the challenge of avoiding immune recognition, but also interact with a multitude of host-associated microbes that influence their ability to successfully colonize the host [28, 50–52]. Our study identifies a link between MHCI and MHCII variability and the gut microbiome of mouse lemurs, and in turn taxa associated with MHC variability predicted parasite infection. Our findings support the notion that natural variability of the MHC is associated to parasite resistance via the microbiome.

Parasite-specific immune responses driven by MHC polymorphisms is extensively supported [53–56], and our study provides evidence of a direct link of the MHC on infection, as suggested by specific MHC motif associations with AdV infection status. We also examined the relative contribution of specific MHC motifs and functional diversity on AdV infection. We found that a specific MHC motif (Migr-DRB*34) is a stronger predictor of AdV infection status in mouse lemurs compared to MHC functional diversity. Nonetheless, we note that MHC functional diversity is more likely to be associated with parasite diversity than infection status of a single parasite [57]. We also found a link between between AdV and helminth infection and distinct aspects of microbial diversity: AdV was associated with the diversity of the microbiome, while helminth infection was related to the divergence of the microbial community. As has been previously reported [30], AdV can have a direct effect on the microbial community. The reciprocal interaction between the hosts’ immune response, gut microbiome and parasites limits our ability to infer the direction of causality using the correlative approach of our study [58]. Nonetheless, our results suggest that the combined role of MHC variability and gut microbiome diversity constitute an important element in explaining parasite infection status.

Given the immunological challenges stemming from high microbial diversity in the gut, we expected that MHC variability would shape microbiome diversity and composition. We found a subtle negative effect of MHCII sequence divergence (i.e. MHCII functional diversity) on the diversity of metabolic pathways (i.e. predicted functional diversity of the microbiome) (r^2^=4.1%). However, we did not find evidence of an effect of MHC diversity at the taxonomic level. Each metabolic pathway is characterised by a set of bacterial taxa (resulting in functional redundancy [59]), suggesting that the effect of MHCII was only observed in pathway-based analyses due to the cumulative effect of combining multiple taxa. We hypothesize that there is a cost to broad antigen-derived immune responses through high MHCII diversity by negatively impacting functional microbial diversity. A cost has previously been suggested through the lens of autoimmune disease [60], whereby there is a trade-off between maximal parasite recognition and the risks associated with autoimmune diseases. Thymic negative selection eliminates T-cells with strong binding affinity to self/MHC complexes [61, 62]. Consequently, high MHC diversity may limit the T-cell receptor repertoire (TCR), leading to suggestions that an optimal intermediate MHC diversity should be selected rather than maximised. However, the trade-off between thymic selection and MHC diversity has been challenged since increased MHC diversity might also enhance positive selection of the TCR repertoire [63]. Thus to date, the mechanisms that constrain MHC diversity remain an enigma [49]. Our recent understanding of microbial diversity within the host requires MHC diversity theory to be revised to account for the potential cost of immunity on the metabolic activities of the microbiome and merits future empirical research.

We evaluated qualitative (specific MHC motifs) and quantitative (MHC functional diversity) MHC effects on gut microbiome composition. We found that both MHC genes were associated with shifts in the composition of the microbial community, with supertype MHCI ST*17 (r^2^ = 9.6%) and MHCII allele Migr-DRB*36 (r^2^= 16.0%) having particularly strong effects. MHCI, but not MHCII, sequence divergence was also linked to among individual differences in microbial communities (r^2^=16.4%). The strong effect of MHCI sequence divergence on microbiome composition contradicted our expectations since MHCII is known to have a higher affinity for extracellular peptides than MHCI. However, in our system, MHCI (median of 7 alleles per individual) was much more diverse than the MHCII (a maximum of 2 alleles per individual), thus among individual variation is much higher in MHCI than MHCII potentially explaining why variation in MHCI was a stronger predictor of microbiome composition than MHCII. Previous work has demonstrated a link between natural variability at MHCII genes and the microbiome [12]. However, the mechanisms through which the MHCI may regulate the gut microbiome remain largely unknown and our results suggests that future research on the role of adaptive immunity in sculpting the gut microbiome should incorporate both MHCI and MHCII molecules.

We hypothesized that specific MHC motifs and high functional diversity would selectively target opportunistic pathogens over commensals [14]. However, we did not find a consistent trend across MHC motifs and functional diversity metrics which predicted both positively and negatively the relative abundance of gut commensals and potentially harmful bacteria. Among the former, for instance, we found an increase in the relative abundance of ASVs assigned to *Odoribacter*, a short-chain fatty acid (SCFA)-producing microbe [44], according to a specific MHCI motif (ST*2). SCFA contributes to the integrity of the epithelial barrier and is an important source of energy [64, 65]. Another SCFA generating microbe, Prevotellaceae UCG-001 [66], was both negatively (ST*12 and Migr-DRB*20) and positively (ST*1) associated to MHCI and MHCII specific motifs. Among potentially harmful bacteria, we observed that a specific MHCII allele (Migr-DRB*36) limited the relative abundance of *Collinsella* and *Campylobacter*, bacteria known to induce inflammatory immune responses by negatively affecting gut permeability [67–69]. In contrast *Campylobacter* was enriched in the presence of the MHCI motif ST*3 and with increasing MHCI functional diversity (MHCI_nST_).

Overall, our results highlight three broad features of the effect of MHC variability on the microbiome. First, both MHCI and MHCII motifs may play a role in selectively sculpting the microbiome. Second, antigen-specific immune responses driven by the MHC are likely to influence the establishment of a few select microbial taxa (commensals and pathogens) rather than overarching control of the whole microbial community supporting previous experimental findings in germ-free mice by Khan et al. [14]. Third, a restricted number of MHC motifs (in the case of mouse lemurs a single MHCII allele, Migr-DRB*36, that is associated with the relative abundance of 22 ASVs) can significantly shape the microbial flora, a result that is in agreement with the strong HLA-disease associations attributed to bacterial pathogenesis in humans [70–72].

A main objective of this study was to evaluate the indirect effect of the MHC on parasite infection via the microbiome. Our data indicates that two taxa are particularly strongly associated with AdV resistance: *Odoribacter*, an SCFA producing microbe, and *Campylobacter*, an opportunistic pathogen. As mentioned above the relative abundance of these microbial biomarkers of AdV resistance is determined by MHC variability. The enrichment of SCFA generating microbes associated with high MHC diversity points at the potential involvement of the immunoregulatory effects of SCFA in promoting tolerance. A key mechanism of T-cell lineage differentiation involves SCFAs which are known to shift the balance towards regulatory Foxp3^+^ T-cells (Treg), limiting inflammation [73–75]. A Treg dominated environment might also be beneficial for opportunistic pathogens, which could explain the observed co-existence between beneficial and potentially harmful bacteria despite high MHC diversity. Pathogenic bacteria can also develop species-specific adaptations to ensure their colonization success [76]. *Campylobacter* uses specialized organelles for adherence and penetration of epithelial cells [77], and *Campylobacter*-specific IgA coating enables *C. jejuni* to aggregate in high densities within the mucus layer, preventing other bacteria from colonizing the gut [78, 79]. There are two possible explanations for the antagonistic interaction between AdV and *Campylobacter* : competitive exclusion and indirect interaction mediated by the immune system. Indeed, both AdV, which is a common persistent infection agent of gut tissue [80], and gut bacterial flora share a common niche. *Campylobacter*-enriched metabolic pathways among AdV^neg^ individuals supports an antagonistic interaction between *Campylobacter* and AdV infection. Alternatively, infection with either of these potential pathogens is likely to lead to inflammatory immune responses, which will have a knock-on effect on the colonization success of non-targeted parasites. Finally, we found that the MHCI supertype ST*3 was a protective motif against AdV (S2 Fig) while also being associated with the enrichment of *Campylobacter* and *Campylobacter*-enriched pathways. The link between AdV and ST*3 might be a result of the effect of this motif on the relative abundance of *Campylobacter* which in turn is negatively associated with AdV infection status.

We did not find a direct link between MHC variability and helminth infection. However, the bacterial community among helminth infected individuals was more similar than the communities among non-infected individuals, a result that is in agreement with previous work demonstrating that by shaping the immunological environment of the gut, helminth infection selects for bacterial taxa tolerant to a Th2 immunological environment [81]. We found an increase in abundance of Prevotellaceae UCG-001, a SCFA generating microbe [66] associated to multiple MHC motifs, among helminth^pos^ individuals, further supporting the potential involvement of the microbiome-helminth interaction promoting tolerance via the expansion of Treg cells. Helminths are also capable of metabolizing SCFA, and it has been previously suggested that a mutualistic interaction between commensals and helminths promotes gut homeostasis [81]. This phenomenon known as the “hygiene-hypothesis” has gained considerable support from epidemiological studies [82, 83] and suggests that helminth infection in mouse lemurs may influence immune responses.

## Conclusion

We demonstrate that the preferential establishment of specific bacterial taxa in a MHC-dependent manner shapes the microbial environment and functional profiles of the bacterial community, potentially influencing the ability of invading parasites to successfully establish an infection. Under this scenario, the host benefits from the antagonistic interactions between specific microbes and parasites, thus providing evidence of an indirect link between MHC variability and parasite resistance via the microbiome. Our understanding of the extraordinary diversity of MHC genes has been focused on parasite-mediated mechanisms, however our findings highlight the potential role of microbiome-driven selection as yet another layer involved in the co-evolutionary dynamics acting on MHC diversity.

## Materials and methods

### Ethics statement

Our work was approved by the ethics committee of the Institute of Zoology of Hamburg University, the University of Antananarivo and Madagascar National Parks. Approval was granted by the Autorisation de Recherche No. 54/13/MEF/SG/DGF/DCB.SAP/SCB of February 22, 2013, issued by the Direction Générale des Forêts and the Direction de la Conservation de la Biodiversité et du Système des Aires Protégées of the Ministère de l’Environnement, et des Forêts and exported to Germany under the CITES permit 576C-EA09/MG14. All animals were handled in accordance with the relevant guidelines and regulations.

### Data collection and parasite screening

We live-trapped *M. griseorufus* at two sites in the Mahafaly Plateau in southwestern Madagascar. Both sites show a semi-arid climate, characterized by irregular rainfall that increases from west to east [84, 85] but differ in the degree of anthropogenic disturbance. While one site, Andranovao (24°01′S; 43°44′E) is located in the Tsimanampetsotsa National Park and consists of intact dry spiny forest, the other one is in Miarintsoa (23°50′S; 44°6′E), a village approx. 40 km east from the National Park consists of open anthropogenic landscape and degraded forest fragments. Sampling was done during the dry seasons of 2013 and 2014 and the wet seasons of 2014 and 2015. We life-trapped *M. griseorufus* using Sherman traps baited with bananas (for details see [86]). After processing of animals and sample collection, we thoroughly cleaned all traps before using them again. Samples were stored at ambient temperature for a few days to weeks in the field, then for variable length of time in a freezer before being transported to Germany where we kept them at −20 C until DNA extraction. We collected small ear biopsies from 143 anesthetized animals and preserved them in 90% ethanol for later MHC characterization. When possible, we collected fecal samples from each trap or handling bag and preserved one subsample in 500 ul RNAlater (Life Technologies) for later adenovirus (AdV) screening (n_Total_= 143; n_Andranovao_= 101 ; n_Miarintsoa_= 42) and microbial sequencing and stored another subsample in 70% ethanol for parasitological analysis (n_Total_= 86; n_Andranovao_= 58 ; n_Miarintsoa_= 28).

### AdV and gastrointestinal parasite screening

We used an Illumina MiSeq platform and followed a target-specific semi-nested PCR assay to evaluate AdV prevalence in 143 mouse lemurs as previously described by Wasimuddin et al. [30]. We counted helminth eggs of a subset of 86 mouse lemurs by using a McMaster flotation technique and a potassium iodide solution [53, 87]. A major concern among epidemological field studies is the low sensitivity of current methods to provide estimates of parasite burden and diversity [88, 89], particularly in wildlife [90]. We therefore use overall helminth infection status as a conservative proxy of gastrointestinal parasite infection. We entered AdV and helminth infection status as presence / absence data for statistical analyses.

### Microbiota sequencing and data processing

Sequencing data of the 16S rRNA gene of the hypervariable V4 region stems from a previous study demonstrating an effect of AdV on the gut microbiome of *M. griseorufus* [30]. The curated dataset used in this study consists of 6,521,596 reads, with a mean coverage of 45,605 reads (min=18950, max=109156) per sample. We processed the microbiome sequencing data with the QIIME2 pipeline [91] and used the implemented DADA2 algorithm [92] for data denoising, merging and calling of amplicon sequence variants (ASVs). For taxonomic assignment we used the Silva database [93]. We excluded all sequences that could not be assigned to a bacterial taxa at the phylum level. We imported and further analysed the microbiome sequencing data output from QIIME2 in the R environment using the PhyloSeq package [94]. We computed microbiome diversity (Faith’s phylogenetic diversity and Shannon’s diversity index) and estimated divergence in microbiome composition (mean distance from the centroid) using Bray-Curtis distance matrix across a 10000 bootstrap sample. For analyses of diversity and composition diversity, we filtered out rare ASVs using a prevalence threshold of <0.1 and <0.2, respectively. Out of the 1505 ASVs identified across the 143 individuals, following prevalence filtering we retained 169 for analyses of diversity and 116 ASVs for analyses of microbial composition.

We predicted functional profiles of the 16S rRNA gene sequencing data at the metabolic pathway level (MetaCyc) by using the PICRUSt2 pipeline with the default parameters [45]. We exported the stratified pathway abundances and taxonomic contributions for analyses in Songbird [40] and in R. The same filtering described above for the taxonomic features was applied for analyses of diversity and composition of the pathway features.

### MHC characterization and diversity estimates

We used high-throughput amplicon sequencing to characterize the MHC class I region (MHCI), MHC class II exon 2 DRB gene (MHCII) on an Illumina MiSeq platform. For MHC characterization, we isolated genomic DNA from ear biopsies by using the Qiagen DNneasy Blood & Tissue Kit (Qiagen). We designed target specific primers to amplify the MHCI gene (fragment length= 236 bp, up to 10 loci; MHCI-Migr-F:

5’-CCCAGGCTCCCACTCCCT-3’and MHCI-Migr-R:

5’-GCGTCGCTCTGGTTGTAGT-3’) and MHCII-DRB gene (fragment length = 171 bp, 1 locus, JS1:5’-GAGTGTCATTTCTACAACGGGACG-3’and JS2

5’-TCCCGTAGTTGTGTCTGCA-3’, [95]. Both MHC genes were previously described as functional regions coding for antigen binding sites [95–97].

We prepared the Illumina sequencing libraries by performing two consecutive rounds of PCR following the approach of Fluidigm System (Access ArrayTM System for Illumina Sequencing Systems; ^©^Fluidigm Corporation). We sequenced the libraries using the Illumina MiSeq platform. Sequencing was done using technical replicates for 85% and 100% of the samples at the MHC class II-DRB and MHC class I gene respectively. Sequence variants of the MHCI gene were identified as true alleles if they were identified in both technical replicates. Negative controls over sequencing runs were clean (< 50 reads after merging). We generated a total of 15,050,630 and 749,428 paired-end reads to characterize allelic diversity at the MHCI genes (average of 51,192 reads, min= 6031, max= 319259) and MHCII gene (average of 2,306 reads, min=392, max=3649), respectively.

Sequence data was processed using the ACACIA pipeline [98] (code available under https://gitlab.com/psc_santos/ACACIA) using a proportion threshold (low-por) of and 0.10 for allele calling on the MHCI and MHCII-DRB dataset, respectively. MHCI is a highly variable region characterized by a series of duplications [96, 97], while the MHCII DRB gene is non-duplicated in mouse lemurs [95, 99, 100]. In contrast to MHCII, the influence of diversity at the MHCI region on parasite resistance remains poorly explored. MHCI and II genes are not linked in mouse lemurs [97] and we therefore expect them to exhibit functional dissimilarities.

We aimed to quantify functional MHC diversity at each MHC gene. To this end we first identified the sites under positive selection (PSS) using the maximum likelihood analysis implemented in codeML (PAML software [101], since these sites are likely to be protein binding sites that recognise antigens. Positive selection is indicated by *d*_N_/*d*_S_ ratio (*ω*) > 1. The following models of codon evolution were computed: M7 (assumes variation of *β* : *ω* among codons modelled under a *β* distribution and does not allow for positive selected sites) and M8 (similar to M7 but assumes *ω* > 1). Model M7 serves as a null model and can be compared to model M8 by means of the likelihood-ratio test. To identify the best fitting model the twice log likelihood difference is compared with a *χ*^2^ distribution. Subsequently, if the model indicating selection (M7) results in a significant better fit to the data, the Bayesian approach in CODEML was used to determine the identity of sites under positive selection. Thirteen and eight positively selected sites were identified in the allele sequences of the MHC I and MHC II genes, respectively (S1 Table). For the MHCI, PSS amino-acid sequences of each allele were described by physicochemical attributes (z-values), and we used a discriminant analysis of principal components (DAPC, R package adegenet [102]) to group MHCI alleles into clusters (supertypes). Given that the MHCII DRB is a non-duplicated gene, we considered functional MHC variants that consist of unique amino-acid sequences (49 out of 51 alleles) at PSS and for statistical analyses, we included alleles present in more than three individuals (frequency > 0.03) (Fig 2). We used the number of MHCI supertypes (MHCI_nST_) per individual as a measure of individual diversity. Since there is only a single locus at the MHCII and 97% of individuals had the maximum of two alleles at this gene, the number of MHCII supertypes was not estimated. In order to assess sequence diversity (divergence of alleles within an individual [103]), we estimated mean sequence divergence over positive selected sites (distPSS) based on the Grantham distance matrix [104] as done in [105] for both the MHCI (MHCI_distPSS_) and the MHCII-DRB (MHCII_distPSS_). This approach enabled us to take into account the physicochemical properties of the PSS amino-acid sequences, being an improved proxy of MHC functional divergence.

### Statistical analyses

We used a probabilistic approach to infer significant associations, (i.e. co-occurrence) between AdV, helminths and MHC functional motifs (i.e. MHCI supertypes and MHCII alleles). We identified the co-occurrence patterns between parasites and MHC diversity using the package “cooccur” [106]. Observed frequencies of parasite-MHC motif pairs that are significantly larger than expected indicate positive associations and significantly lower than expected indicate negative associations. Were corrected *P*-values for multiple comparisons across MHC motifs using the Bonferroni correction. MHC motifs that were significantly associated with parasite infection were included as predictor variables for downstream analyses. The inclusion of MHC motifs along with MHC functional diversity allowed us to estimate the relative contribution of specific alleles (i.e. qualitative MHC effects) vs functional diversity (i.e. quantitative MHC effects), providing insights on the distinct selective mechanisms of the MHC operating on parasite infection and microbiome diversity and composition [107].

To assess the link of parameters on infection status, we fitted GLMs with a binomial error structure and ranked them using the information-theoretic (I-T) model selection procedures, with the Akaike information criterion adjusted for small sample sizes (AICc) with the MuMIn package [108]. We used model averaging (without shrinkage) across the credible set of models (cumulative AICc weight < 0.95) to generate parameter estimates. We interpreted the effect of variables on our response variable if 95% confidence intervals did not overlap with zero. We report effect sizes using partial-r for continuous variables and Cohen’s D for categorical variables and estimate 95% confidence intervals by boostrap (n = 10000) [109].

We fitted linear term for MHCI_nST_, MHCI_distPSS_, and MHCII_distPSS_ and covariates in a GLM with a gamma distribution to test for the association between MHC functional diversity and alpha diversity of the microbiome. We used model selection and 95% confidence intervals to interpret the effect of these predictors on microbiome alpha diversity.

We explored the contribution of the predictor variables on the bacterial (taxa) and functional (metabolic pathways) community structure by using canonical correspondence analysis (CCA) using the vegan package in R [110] on a chord transformed matrix. We tested for significance of the overall CCA model by means of permutation (9999). We used the *envfit()* function to assess the significance of the fitted vectors (MHC functional diversity) and factors (MHC specific motifs, site, season, sex, AdV and helminth infection). The effect of each MHC motifs was assessed independently in separate models along with all other covariates.

To explore the effects of MHC class I and class II motifs on the relative differential abundance of specific ASVs, we applied an analysis of composition of microbes (ANCOM) [111] which controlled for the effect of site and season. ANCOM runs a set of pairwise tests for each ASVi and across each ASVj, where the null sub-hypothesis is that the log ratio ASVi/ASVj is not associated with a given predictor (the total number of sub-hypothesis is the total number of ASVs investigated minus one). A W score is the sum of rejected null sub-hypotheses. Therefore, W estimates the strength of support for a relative differential abundance of a specific ASV according to a predictor of interest. We used volcano plots to visualize the relationship between the W score and the estimates of the differential logs from a linear model. We used a W score threshold of 0.6, which is equivalent to 60% null sub-hypotheses rejected across the set of pairwise tests, and a *P* value < 0.005 to identify ASVs whose relative abundance differs a according to the presence of specific MHC motif. We used differential ranking [40] to identify ASVs most associated with low, and high MHC diversity by means of a multinomial regression model with MHCI_distPSS_, MHCI_nST_ and MHCII_distPSS_, site and season as covariates, using Songbird [40]. We used a batch size of 5, epochs of 50,000 and a prior of 0.8 as parameters for the analysis. The coefficients of the multinomial regression represent the log fold change for each ASV along the range of MHC functional diversity estimates.

In order to investigate whether microbial taxa associated to specific MHC motifs and MHC functional diversity contribute to AdV and helminth infection status, we used a reference frame approach. Traditional methods of compositional analyses do not allow to make interpretations of absolute differences in ASV abundance. However, by using a carefully chosen reference that is not expected to differ in absolute loads across covariates and is known to be stable across environments, the reference frame allows to overcome the limitations associated with interpreting differences in relative abundance and to make some cautious inference about absolute differences [40]. To calculate log-ratios, we defined *Bifidobacterium* spp. as a reference frame given that: i) exhibit high prevalence and, ii) we expect this taxon to be relatively stable within the microbial community because a series of strategies that enable host tolerance have been identified among species of bifidobacteria. Indeed, previous studies have demonstrated that immuno-modulatory roles of bifidobacteria can lead to reduced inflammation [112, 113] and the production of key molecules enable bifidobacteria to withstand stressful environmental conditions within the gut [18, 114], making this taxon a useful reference of health. Here we report the differential abundance of taxa according to AdV and helminth infection relative to the abundance of *Bifidobacterium* spp. using a Mann-Whitney Wilcoxon test. We considered common microbial taxa (with a prevalence *>* 60%) that were ranked high or low along this gradient as suitable candidates for assessing (with a log-ratio test) if infection status influences their relative abundance. We controlled for multiple comparisons across taxa by means of Bonferroni correction of *P*-values. We report estimates of effect size (Cohen’s D and 95% confidence intervals) and interpret effects > |0.50|.

We also used ANCOM and differential ranking to identify the metabolic pathways associated with MHC motifs and functional diversity, respectively, following the same approach used for taxonomic based analyses. We then used the subset of pathways linked to MHC variability to test whether they predict AdV and helminth infection status. We used with MHCI_distPSS_, MHCI_nST_ and MHCII_distPSS_, site, and season as covariates in the multinomial regression model with a batch size of 5, epochs of 50,000 and a prior of 1 as parameters for the analysis using Songbird. To calculate log-ratios we used the TCA pathway (PWY-7254) as a reference since this is a common pathway central to the generation of energy and present in most aerobic living organisms.

## Acknowledgments

The study was carried out under the partnership agreement between MNP (Madagascar National Parks), the Department of Animal Biology, University of Antananarivo and the Department of Animal Ecology and Conservation, University of Hamburg. This study was funded by SuLaMa/BMBF (FKZ 01LL0914) and German Science Foundation, DFG, SPP 1596 “Ecology and Species Barriers in Emerging Viral Diseases”, Ga 342/19-1. We appreciate the help and the logistic support provided by all the field assistants, and Kerstin Wilhelm and Ulrike Stehle for their support in the lab.

## Supporting information

**S1 Fig. Clustering of MHC class I alleles into supertypes.** Clustering of MHC class I alleles into supertypes using discriminant analysis of principle components (DAPC) based on a matrix of physiochemical properties of the sites under positive selection (PSS) of each allele. *(A)* Number of clusters chosen using the find.clusters() function of the adegenet package [102]. The red dashed line shows *(k)*= 17. *(B)* Scatterplot of the first and second discriminant functions showing the supertype clusters.

**S2 Fig. MHC class II DRB allele frequency.** Relative frequencies of MHC class II DRB alleles in *Microcebus griseorufus* (n=143). The identity of allele variants sharing identical amino–acid sequences at positive selected sites (PSS) is shown in parenthesis.

**S3 Fig. Association between specific MHCI and MHCII motifs and AdV and helminth infection status.** Relative motif frequencies of MHCI supertypes *(A)* and MHCII alleles found in more than 3 individuals *(B)*. *(C-D)* Expected and observed co–occurrence probabilities between MHCI supertypes and AdV and helminth infection status. *(E-F)* Expected and observed co-occurrence probabilities between MHCII alleles and AdV and helminth infection status. Asterisks indicate statistical significance between expected and observed parasite infection, based on co occurrence analysis. * *P* < 0.05; ** *P* < 0.005.

**S4 Fig. Canonical correspondence analysis (CCA) biplot for taxonomic composition in association with MHC functional diversity.** Canonical correspondence analysis (CCA) for taxonomic composition in association with MHC functional diversity (n=143). CCA biplot depicting ASVs (full circles) and MHC diversity estimates as arrows. MHCI_distPSS_ (r^2^=0.16, *P*<0.0001) functional diversity significantly predicting microbiome composition is shown in red. Labels correspond to assigned taxa.

**S5 Fig. CCA biplot depicting the differences among samples according to MHCI supertypes.** CCA biplot depicting the differences among samples according to the presence/absence of the MHCI supertypes.

**S6 Fig. CCA biplot depicting the differences among samples according to MHCII alleles.** CCA biplot depicting the differences among samples according to the presence/absence of the MHCII alleles.

**S7 Fig. CCA biplot depicting the differences among samples according to AdV and helminth infection status and covariates.** CCA biplot depicting the differences among samples according to AdV and helminth infection status and covariates.

**S8 Fig. CCA biplot depicting the differences among samples according to MHCI and MHCII motifs.** CCA biplot depicting the differences among samples according to the presence/absence of the MHCI supertypes and MHCII alleles.

**S9 Fig. The effect of covariates on functional pathway diversity and composition.** Boxplots illustrating differences in pathway eveness according to season *(A)*, pathway richness according to site*(B)* and CCA biplot depicting differences among samples according to season *(C)*.

**S1 Table. Polymorphic sites exhibiting signals of positive selection.**

Log-likelihood values and parameter estimates of models testing for positive selection acting on MHC class I and MHC class II DRB exon 2 of *Microcebus griseorufus*. Parameters *p* and *q* computed from the beta distribution. *ω*= d_N_/d_S_ ratio. p_n_= of sites that fall into *ω*n site class. Site positions inferred to be under positive selection estimated at a * 95% and ** 99% confidence interval level.

**S2 Table. Top ranking GLMs examining the association between MHC, microbiome diversity (Faith’s PD), and covariates on AdV infection status.** Model selection of GLMs (top 20 models subset) examining the association between MHCI and MHCII motifs and diversity estimates, microbiome diversity (Faiths’ PD) and divergence (Bray-Curtis) and covariates on AdV infection status. AIC_c_= Akaike Information Criterion for small sample sizes; AIC_c_*ω* = AIC weight; “+” denotes categorical parameters included in the model whilst the inclusion of a continuous variables is indicated with its estimate.

**S3 Table. Top ranking GLMs examining the association between MHC, microbiome diversity (Faith’s PD) and divergence, and covariates on helminth infection status.** Model selection of GLMs (top 20 models subset) examining the association between MHCI and MHCII motifs and diversity estimates, microbiome diversity (Faiths’ PD) and divergence (Bray-Curtis) and covariates on helminth infection probability. AIC_c_= Akaike Information Criterion for small sample sizes; AIC_c_*ω* = AIC weight; “+” denotes categorical parameters included in the model whilst the inclusion of a continuous variables is indicated with its estimate.

**S4 Table. Top ranking GLMs examining the association between MHC, microbiome diversity (Shannon’s index), and covariates on AdV infection status.** Model selection of GLMs (top 20 models subset) examining the association between MHCI and MHCII motifs and diversity estimates, microbiome diversity (Shannon’s index) and divergence (Bray-Curtis) and covariates on AdV infection status. AIC_c_= Akaike Information Criterion for small sample sizes; AIC_c_*ω* = AIC weight; “+” denotes categorical parameters included in the model whilst the inclusion of a continuous variables is indicated with its estimate.

**S5 Table. Top ranking GLMs examining the association between MHC, microbiome diversity (Shannon’s index), and covariates on helminth infection status.** Model selection of GLMs (top 20 models subset) examining the association between MHCI and MHCII motifs and diversity estimates, microbiome diversity (Shannon’s index) and divergence (Bray-Curtis) and covariates on helminth infection probability. AIC_c_= Akaike Information Criterion for small sample sizes; AIC_c_*ω* = AIC weight; “+” denotes categorical parameters included in the model whilst the inclusion of a continuous variables is indicated with its estimate.

**S6 Table. Model average parameter estimates and 95% confidence intervals of GLMs examining the association between MHCI and MHCII diversity estimates, microbiome diversity (Faiths’ PD) and divergence (Bray–Curtis) and covariates on AdV and helminth infection status.**

**S7 Table. Model average parameter estimates and 95% confidence intervals of GLMs examining the association between MHCI and MHCII diversity estimates, microbiome diversity (Shannon’s index) and divergence (Bray–Curtis) and covariates on AdV and helminth infection status.**

**S8 Table. Model average parameter estimates and 95% confidence intervals of GLMs examining the association between phylogenetic diversity (Faiths’ PD) and evenness (Shannon’s diversity index), MHCI and MHCII functional diversity, and covariates.**

**S9 Table. Model average parameter estimates and 95% confidence intervals of GLMs examining the association between predicted pathway richness and evenness (Faith’s PD), MHCI and MHCII functional diversity, AdV infection status, and covariates.**

**S10 Table. Model average parameter estimates and 95% confidence intervals of GLMs examining the association between predicted pathway richness and evenness (Shannon’s diversity index), MHCI and MHCII functional diversity, helminth infection status, and covariates.**

